# Altered Metabolism During the Dark Period in *Drosophila* Short Sleep Mutants

**DOI:** 10.1101/2023.10.30.564858

**Authors:** Dania M. Malik, Arjun Sengupta, Amita Sehgal, Aalim M. Weljie

## Abstract

Sleep is an almost universally required state in biology. Disrupted sleep has been associated with adverse health risks including metabolic perturbations. Sleep is in part regulated via circadian mechanisms, however, metabolic dysfunction at different times of day arising from sleep disruption is unclear. We used targeted liquid chromatography-mass spectrometry to probe metabolic alterations using high-resolution temporal sampling of two *Drosophila* short sleep mutants, *fumin* and *sleepless*, across a circadian day. Discriminant analyses revealed overall distinct metabolic profiles for mutants when compared to a wild type dataset. Altered levels of metabolites involved in nicotinate/nicotinamide, alanine, aspartate, and glutamate, glyoxylate and dicarboxylate metabolism, and the TCA cycle were observed in mutants suggesting increased energetic demands. Furthermore, rhythmicity analyses revealed fewer 24 hr rhythmic metabolites in both mutants. Interestingly, mutants displayed two major peaks in phases while wild type displayed phases that were less concerted. In contrast to 24 hr rhythmic metabolites, an increase in the number of 12 hr rhythmic metabolites was observed in *fumin* while *sleepless* displayed a decrease. These results support that decreased sleep alters the overall metabolic profile with short sleep mutants displaying altered metabolite levels associated with a number of pathways in addition to altered neurotransmitter levels.

## Introduction

Despite being an essential process for most organisms, the role of sleep is not well understood. The state of sleep is a tightly controlled process in part by circadian regulation and homeostatic pressures [1–3]. Among several proposed purposes of sleep, such as facilitating the clearance of toxins and memory formation, it is thought to play a role in overall metabolic restoration [4–6]. Disrupted sleep has not only been identified as a risk factor for pathophysiolgies, such as neurodegenerative disorders, but has also been associated with altered metabolism in both humans and model organisms [7, 8]. Given the potential role sleep plays in metabolic restoration and the role circadian processes play in its regulation, it reasons that metabolic rhythmicity may facilitate separation of incompatible processes. Altered parameters associated with cycling such as phases and amplitudes have been noted as a result of sleep disruption [9, 10]. However, not all sleep studies have used high time resolution sampling and thus provide limiting insights on cyclical processes and time of day variability.

Although the role of the clock and environmental factors has been explored in *Drosophila*, a comprehensive screen using high time sampling in short sleep models has not been performed. We sought to identify sleep-driven changes as a function of sleep as well as time of day in *Drosophila*. *Drosophila* enables the study of both circadian and sleep processes simultaneously as it not only has a molecular clock similar to the mammalian one but also enters a state of inactivity which meets the criteria for sleep [3, 7, 11, 12]. A number of short sleep mutants have been identified in *Drosophila* [13–17]. We incorporated two short sleep mutants, *fumin (fmn)* [13]. and *sleepless (sss)* [14], as chronic models of sleep loss in order to be able to differentiate possible sleep driven changes from those specific to the mutants. Decreased sleep in *fmn* arises from a dopamine transporter defect resulting in impaired dopamine reuptake at the synapse while although the mechanism in *sss* is unclear, the loss of the *sleepless* protein is thought to play a role in membrane excitability and synaptic transmission and results in decreased GABA levels and decreased sleep [13, 14, 18–20]. Since both mutants have decreased sleep arising from different mutations, together they can begin to provide a unique metabolic fingerprint of decreased sleep. We performed a steady state metabolomics analysis on both sleep mutants at 2-hour resolution and compared with a previously collected wild type data set. Through this, common metabolome-level changes were observed across the short sleep mutants with overall altered nicotinate and nicotinamide metabolism. Additional metabolic pathways were observed to be impacted during the dark period with additional differences noted in rhythmicity.

## Results

### Analytical quality is robust across datasets

In order to identify and separate sleep-driven changes in *Drosophila*, we incorporated the sleep mutant dataset with a previously acquired dataset under similar conditions containing wild type (WT, isogenic controls) samples under light:dark (LD) conditions [21]. We reintegrated the WT dataset alongside the sleep mutant data in order to minimize any variability arising from inter individual differences in identifying peaks and to ensure the same peak was integrated in a similar method across all samples (Figure 1A). The ion counts from each dataset were independently corrected for instrument drift and normalized using the same procedures as described in the methods. Principal component analysis (PCA) was used to assess the data quality of each dataset independently. Clustering of quality control (QC) samples and evenly distributed biological samples were noted in the scores plot while the corresponding loadings plots also showed evenly distributed metabolites (Figure 1B and C). Taken together, this supports robust analytical quality in each dataset. The datasets were then merged based on identified metabolite peaks and technical replicates were averaged. Since the datasets were acquired at different times, to minimize the impact of batch differences, we chose to normalize the corrected ion counts for each time point to the corrected ion counts at ZT 0 (lights on). This was performed on a per metabolite basis within each biological replicate for each genotype. The resulting PCA also exhibited clustering of the QC samples and evenly distributed samples across the two datasets in the scores plot and evenly distributed metabolites in the loadings plot (Figure 1D). This ZT 0 normalized dataset was used for further analysis as no notable major differences were observed as a function of different collection times.

**Figure 1.**
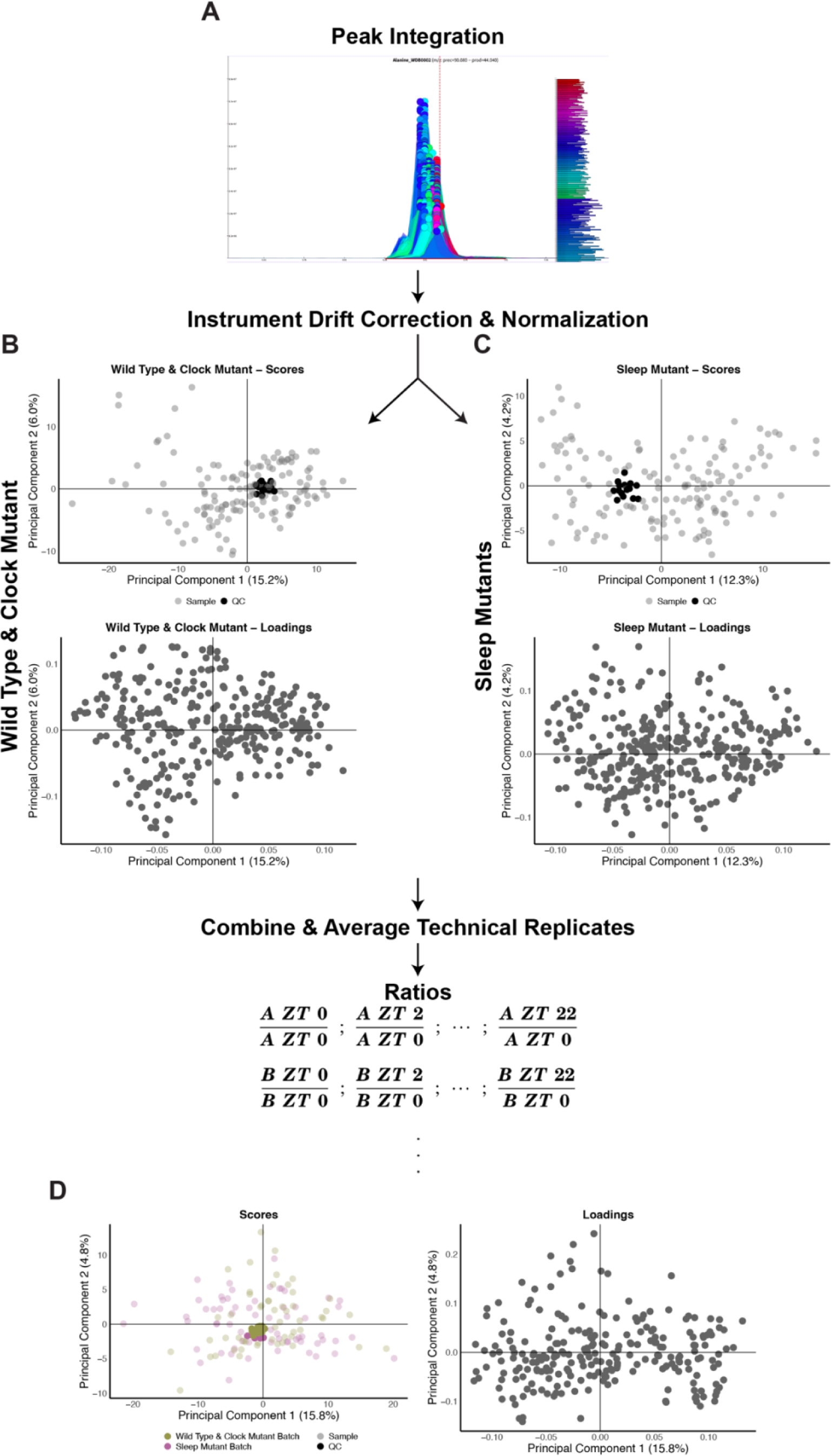
Analytical quality is robust across datasets. (A to D) Simplified overview of data processing and normalization procedure. (A) Example of a chromatographic peak integrated in El-Maven across all samples. (B to C) PCA Scores (top panel) and loadings (bottom panel) plots for LOESS corrected and median fold change (MFC) normalized datasets. Shading in scores plots represent samples (lighter) and QC samples (darker). (D) PCA Scores (left panel) and loadings (right panel) for the combined data after ratios were calculated. Colors represent the two datasets with shading representing samples (lighter shade) and QC samples (darker shade).

### Unique and conserved metabolome differences are present in the short sleep mutants compared to wild type

The two short sleep mutants exhibit reduced sleep due to different mutations. Therefore, to understand whether they also had overall metabolome differences, we employed a multivariate approach using an orthogonal partial least squares-discriminant analysis (OPLS-DA). Here, a significant model was generated when the three genotypes (WT, *fumin,* and *sleepless*) were defined as classes (Figure 2A). The corresponding loadings plot also highlighted metabolites that were significantly contributing to the differences across the three genotypes through variable importance on projection (VIP) values greater than one (Figure 2 B). Together, the scores and loadings plot indicated the presence of overall metabolome-level differences in the genotypes.

**Figure 2.**
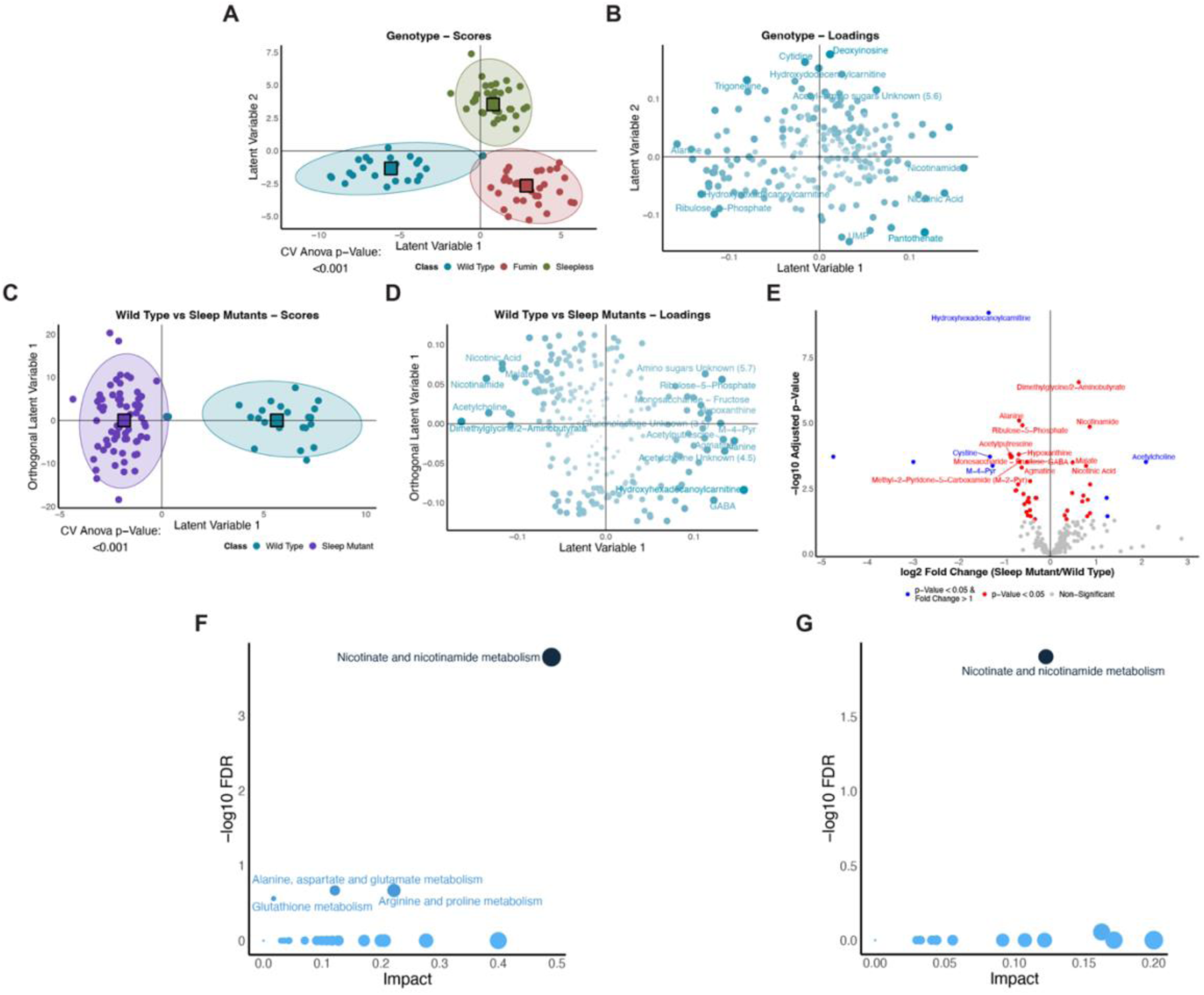
Unique and conserved metabolome differences are present in short sleep mutants in comparison to wild type. (A and C) Scores plots for significant genotype (A) and Wild Type vs Sleep Mutants (C) OPLS-DA models. Colors represent classes. Circle points represent individual samples and square points represent the class average. (A) Classes were defined by genotype. (C) Two classes were defined as Wild Type and Sleep Mutant (*fumin* and *sleepless*). (B and D) Corresponding loadings plots for the significant genotype (A) and Wild Type vs Sleep Mutants (C) OPLS-DA models. Shading and size represent significance as defined by VIP values with larger and darker points representing a higher VIP value. Metabolites with a VIP > 1.5 are labelled. (E) Volcano plot of metabolites analyzed using univariate analyses. Log2 of the fold change between the sleep mutants and wild type is shown on the x-axis and the negative log10 BH adjusted group p-value from a two-way ANOVA is shown on the y-axis. Significance was defined as an adjusted p-value less than 0.05 (red) and adjusted p-value less than 0.05 with a log2 fold change greater than 1 (blue). (F and G) Overview of pathway analysis using VIP metabolites (F, metabolites identified in D) and significant metabolites (E, metabolites identified in E) depicting the negative log10 of the FDR and impact for pathways. Pathways with an FDR less than 0.3 and impact greater than one are labelled. Shading represents significance as defined by FDR values with darker representing smaller FDR values. Size of points represents impact.

However, as we were interested in sleep-driven changes, we sought to determine common differences in the short sleep mutants from the wild type samples. Here, we used an approach of defining two classes where WT samples comprised the first class and the second class consisted of the two short sleep mutants, *fmn* and *sss*. By assigning both short sleep mutants to a single class, the resulting significant metabolites as defined using variable importance of projection (VIP) values would highlight conserved changes occurring in the same direction in both short sleep mutants in contrast to WT. For example, a given metabolite would need to be either increased or decreased in both sleep mutants rather than being increased in one and decreased in the other. Overall, using all time points, a significant discriminant model was generated and able to separate out the sleep mutants defined as a class from the wild type samples (Figure 2C). The corresponding loadings plot also highlighted differences in metabolite levels across the two groups (Figure 2D). For example, higher levels of metabolites such as fructose and ribulose-5-phosphate were associated with the wild type samples while elevated levels of nicotinic acid, malate and acetylcholine were associated with both short sleep mutants. Metabolic pathway analysis was then performed to determine whether the metabolites significantly contributing to the difference of WT from the short sleep mutant samples (metabolites with a VIP value greater than one) were localized to particular metabolic functions. Nicotinate and nicotinamide metabolism was found to be enriched (FDR < 0.2) while pathways such as alanine, aspartate, and glutamate metabolism, arginine and proline metabolism, and glutathione metabolism were also likely implicated with a higher possibility of false discovery (FDR < 0.3) (Figure 2F). A univariate approach using genotype-level FDR adjusted p-values from a two-way ANOVA to identifying differences also highlighted similar changes (Figure 2E). For example, acetylcholine levels were at least twice as high in the short sleep mutants than wild type samples while metabolites such as nicotinamide, nicotinic acid, malate and dimethylglycine/2-aminobutyrate were significantly (p-value less than 0.05) higher in the short sleep mutants as well although did not meet the fold change cut-off of 1. These metabolites were also VIP metabolites in the multivariate analysis and higher levels were associated with the short sleep mutants. Similarly, for wild type, metabolites such as hydroxyhexadecanoylcarnitine and N1-Methyl-4-pyridone-3-carboxamide (M-4-Pyr) levels were significantly higher in WT both through univariate (at least twice as high in WT compared to the short sleep mutants) and discriminant analysis (VIP greater than 1). Other metabolites such as alanine, ribulose-5-phosphate, GABA, and hypoxanthine were also significantly different and associated with higher levels in WT through both analyses. Similar to the pathway results using the VIP metabolites, nicotinate and nicotinamide metabolism also emerged as a significantly enriched pathway when significant metabolites (BH adjusted group p-value from a two-way ANOVA less than 0.05) were used in a pathway analysis (Figure 2G). Overall, the short sleep mutants display both unique and conserved metabolome-level differences and altered nicotinate and nicotinamide metabolism.

### Multiple metabolic pathways are impacted in short sleep mutants during the dark period

Since sleep can be influenced by circadian processes, we next wanted to examine whether the short sleep mutants displayed differences based on time of day. The WT and short sleep mutant samples clustered separately in significant discriminant models for both the light and dark time points (Figure 3A and B). The corresponding loadings plots highlighted metabolites associated with each group during the specific times of day. For example, during the light period, the short sleep mutants were associated with higher levels of metabolites such as acetylcholine while wild type samples were associated with elevated levels of metabolites such as gluconate and acetylputrescine (Figure 4A). On the other hand, during the dark time points, metabolites such as carbamoyl phosphates, nicotinic acid, and nicotinamide were elevated in the short sleep mutants while metabolites including alanine, GABA, and agmatine were elevated in wild type samples (Figure 4D). To further assess time of day differences, we tested whether time of day differences persisted either during the early (first two time points) or late (last two time points) time points during the light or dark period. All comparisons (early light, late light, early dark, and late dark time points) between WT and short sleep mutants resulted in significant discriminant models (Figures 3C to F). The corresponding loadings plots also indicated differences in metabolites associated with each group during the different time periods tested (Figures 4B to C and E to F). Overall, this indicated that the short sleep mutants were distinct from wild type samples in their metabolite profiles not only during the overall light and dark periods but also during shorter blocks of time. Next, to determine whether significantly contributing metabolites were enriched in similar pathways at the different times, we performed a pathway analysis for each of the time periods. Overall, the dark time period generally was associated with a greater number of significantly enriched pathways (Figure 3 G). Three general groups were noted in the significant pathway hits: 1. Pathways were significant overall between the groups irrespective of time and during the overall light and dark period such as nicotinate and nicotinamide metabolism; 2. Pathways were not significant overall but were significant during the overall dark period such as alanine, aspartate, and glutamate metabolism; and 3. Pathways that were not significant during the overall light or dark time periods but were significant when looking at the early and/or late periods such as arginine biosynthesis (late light), glyoxylate and dicarboxylate metabolism (late dark), and the TCA cycle (late dark) (Figure 3G). Taken together, these results point to overall differences in the metabolome of short sleep mutants which may be driven by decreased sleep, however highlight a greater impact on metabolic pathways during the dark time period.

**Figure 3.**
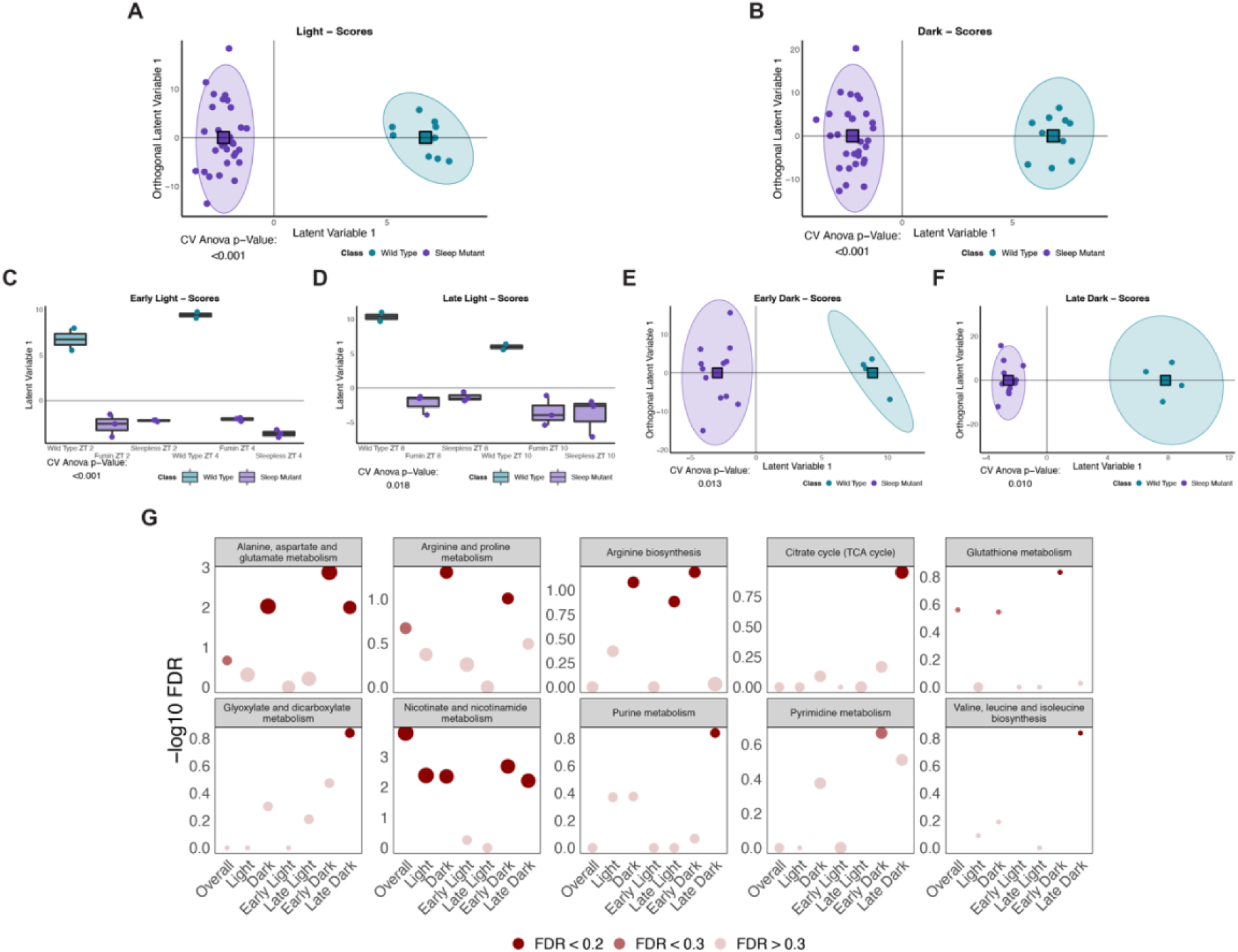
Multiple metabolic pathways are impacted in short sleep mutants during the dark period. (A to F) Scores plots for significant discriminant analysis models. Colors represent classes with circle points representing individual samples and square points representing class averages. Classes were defined as wild type and sleep mutant (*fumin* and *sleepless*) using the following time points: (A) Light time points (ZT 2, ZT 4, ZT 6, ZT 8, and ZT 10); (B) Dark time points (ZT 14, ZT 16, ZT 18, ZT 20, and ZT 22); (C) Early light time points (ZT 2 and ZT 4); (D) Late light time points (ZT 8 and ZT 10); (E) Early dark time points (ZT 14 and ZT 16); and (F) Late dark time points (ZT 20 and 22). (G) Negative log10 FDR values for pathways significant in at least one OPLS-DA model (Figure 2F and A to F). Size represents impact and significance is indicated by shading with a FDR less than 0.2 represented by dark points, FDR between 0.2 and 0.3 represented by medium-shaded points and FDR greater than 0.3 represented by lightest points.

**Figure 4.**
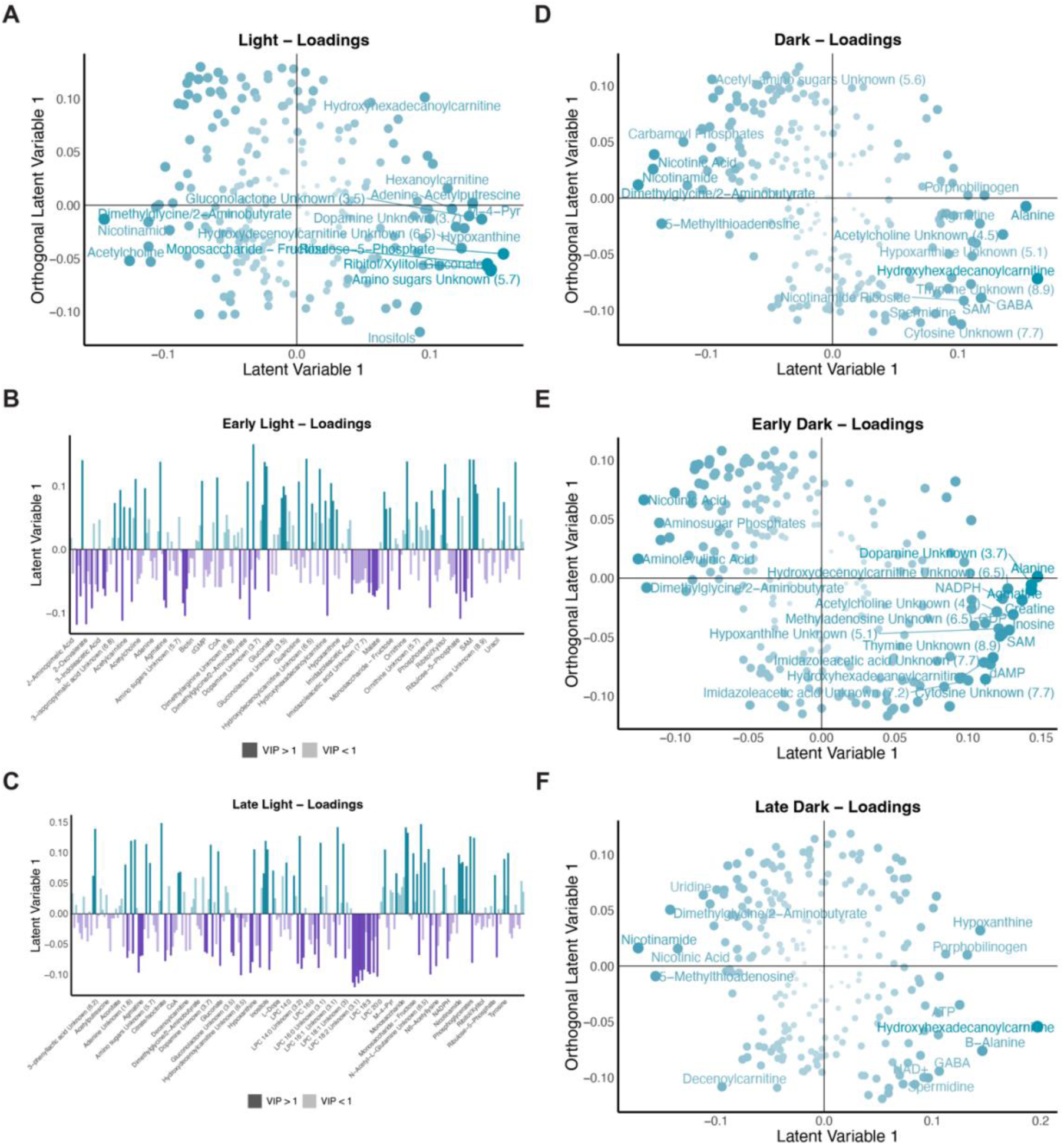
Overview of significant metabolite changes in short sleep mutants. (A to F) Corresponding loadings plot for scores plot (Figure 3). (A, D, E and F) Shading and size represents significance as defined by VIP values with larger and darker points representing higher VIP values. Metabolites with a VIP greater than 1.5 are labelled. (B and C) Colors represent the class associated with higher levels of the metabolite. Shading represents VIP significance with dark indicating a VIP value greater than one. Metabolite names are indicated for those with a VIP greater than 1.5. Loadings plots correspond to the following Wild Type vs Sleep Mutant OPLS-DA models: (A) Light time points (Figure 3A); (B) Early light time points (Figure 3C); (C) Late light time points (Figure 3D); (D) Dark time points (Figure 3B); (E) Early dark time points (Figure 3E); and (F) Late dark time points (Figure 3F).

### Nicotinate and nicotinamide metabolism is affected in short sleep mutants throughout the day

Nicotinate and nicotinamide metabolism was observed to be a significantly enriched pathway across all discriminant models generated except during the early light and late light time periods (Figure 5A). We next looked at the whether the VIP metabolites included in this pathway across the different discriminant models were also significantly impacted at the univariate level between the different groups both in a similar way and during the times when the pathway was significantly enriched. Overall, metabolites associated with this pathway were significantly altered in the short sleep mutants together irrespective of time with metabolites such as NAD+, NADP+, nicotinamide riboside and nicotinamide ribotide displaying decreased levels in the short sleep mutants while increased levels of nicotinamide and nicotinic acid were noted (Figures 5B to G). Interestingly, all of the metabolites except NAD+ were significantly altered during the dark period in the short sleep mutants with similar directionality. Significant differences between wild type and the short sleep mutants were also noted during both the early and late dark periods for nicotinamide and nicotinic acid while nicotinamide riboside and nicotinamide ribotide were only significant during the early dark periods. On the other hand, significant differences during the overall light period were noted for nicotinamide and nicotinamide ribotide between wild type and the two short sleep mutants together (Figures 5D and F). However, across all of the time periods, trends in the directionality of the changes were observed to be consistent with the changes noted earlier between the groups irrespective of time. Taken together, this highlights the impact sleep can have on nicotinate and nicotinamide metabolism across the day.

**Figure 5.**
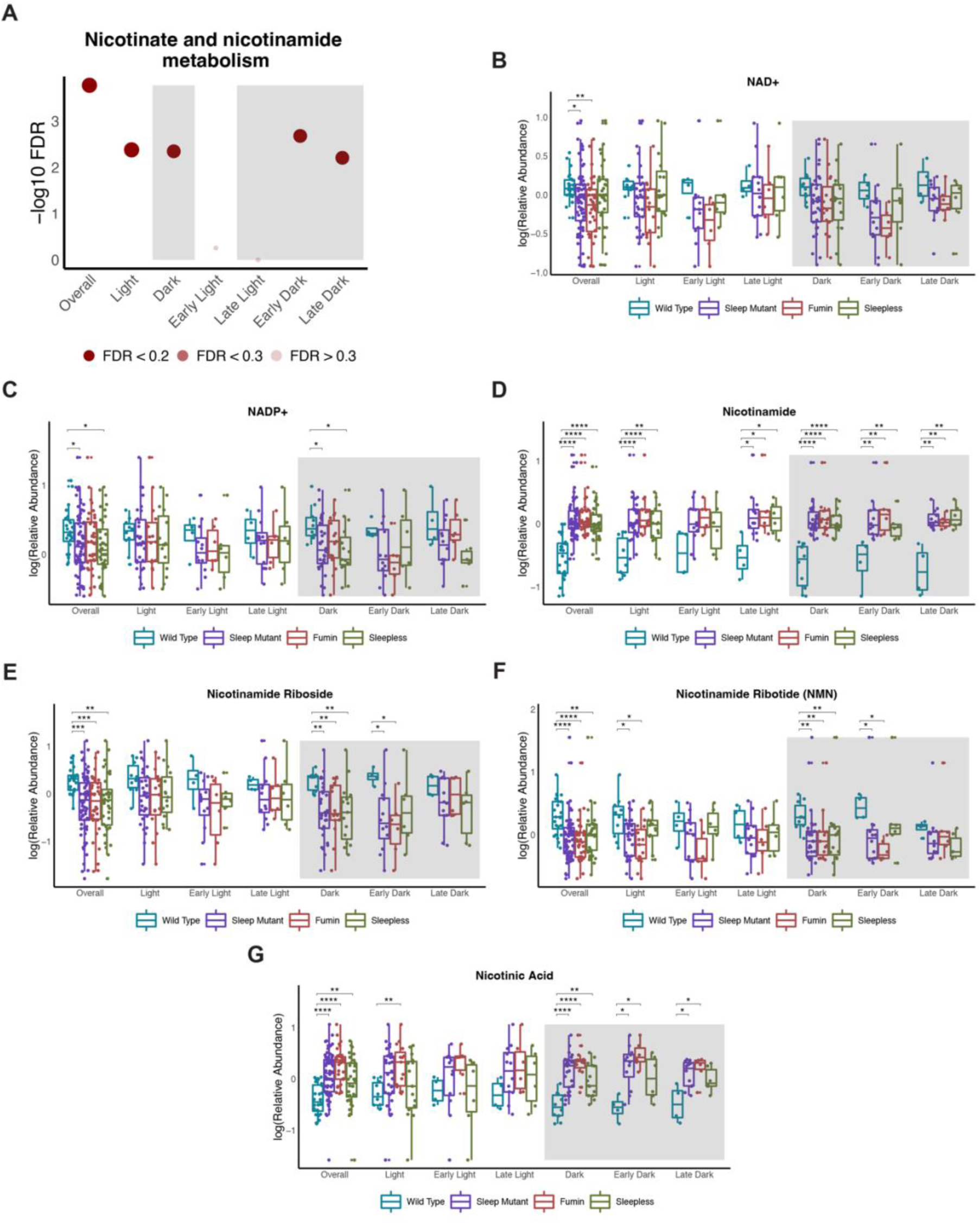
Nicotinate and nicotinamide metabolism is affected in short sleep mutants throughout the day. (A) Negative log10 FDR values for nicotinate and nicotinamide metabolism pathway across the different significant OPLS-DA models. Shading represents FDR values. (B to G) Box plots of metabolites associated with nicotinate and nicotinamide metabolism by groups (wild type, sleep mutants (consists of *fumin* and *sleepless)*, *fumin*, and *sleepless*) shown for different time periods. Significance was defined as a BH adjusted p-value less than 0.05 from a Wilcoxon test within each time period. Significance is indicated as follows: * < 0.05, ** < 0.01, *** < 0.001, **** < 0.0001.

### Arginine related metabolism is altered during the dark time period

Arginine and proline metabolism was also identified as potentially important from the VIP-significant metabolites from the overall WT vs Sleep Mutant OPLS-DA model was (Figure 6 A). This pathway was also significant during the overall dark and early dark time points. The related arginine biosynthesis pathway was also noted to be significant during the overall dark, late light and early dark time points (Figure 6G). In general, metabolites associated with arginine and proline metabolism were observed to be significant between wild type and the short sleep mutants irrespective of time (Figures 6B to F) while metabolites associated with arginine biosynthesis such as citrulline were not significant (Figure 6H). Similarly, arginine and proline metabolism related metabolites such as proline were significantly increased while SAM and spermidine were significantly decreased in the short sleep mutant when compared to wild type during the dark period (Figures 6C, D and E). Even though the overall pathway was noted to be significant during the early dark period, only SAM was significantly decreased during the early dark time points (Figure 6D). Other metabolite levels were trending towards altered levels in the short sleep mutants compared to wild type samples. Furthermore, metabolites related to arginine biosynthesis in general did not show significant changes when measured through a univariate approach (Figure 6H). Overall, both analyses indicated similar overall trends, although differences in metabolites reaching significance was observed. This is however to be expected as a multivariate approach is designed to detect concerted changes across metabolites while a univariate approach looks to determine whether a given metabolite is altered.

**Figure 6.**
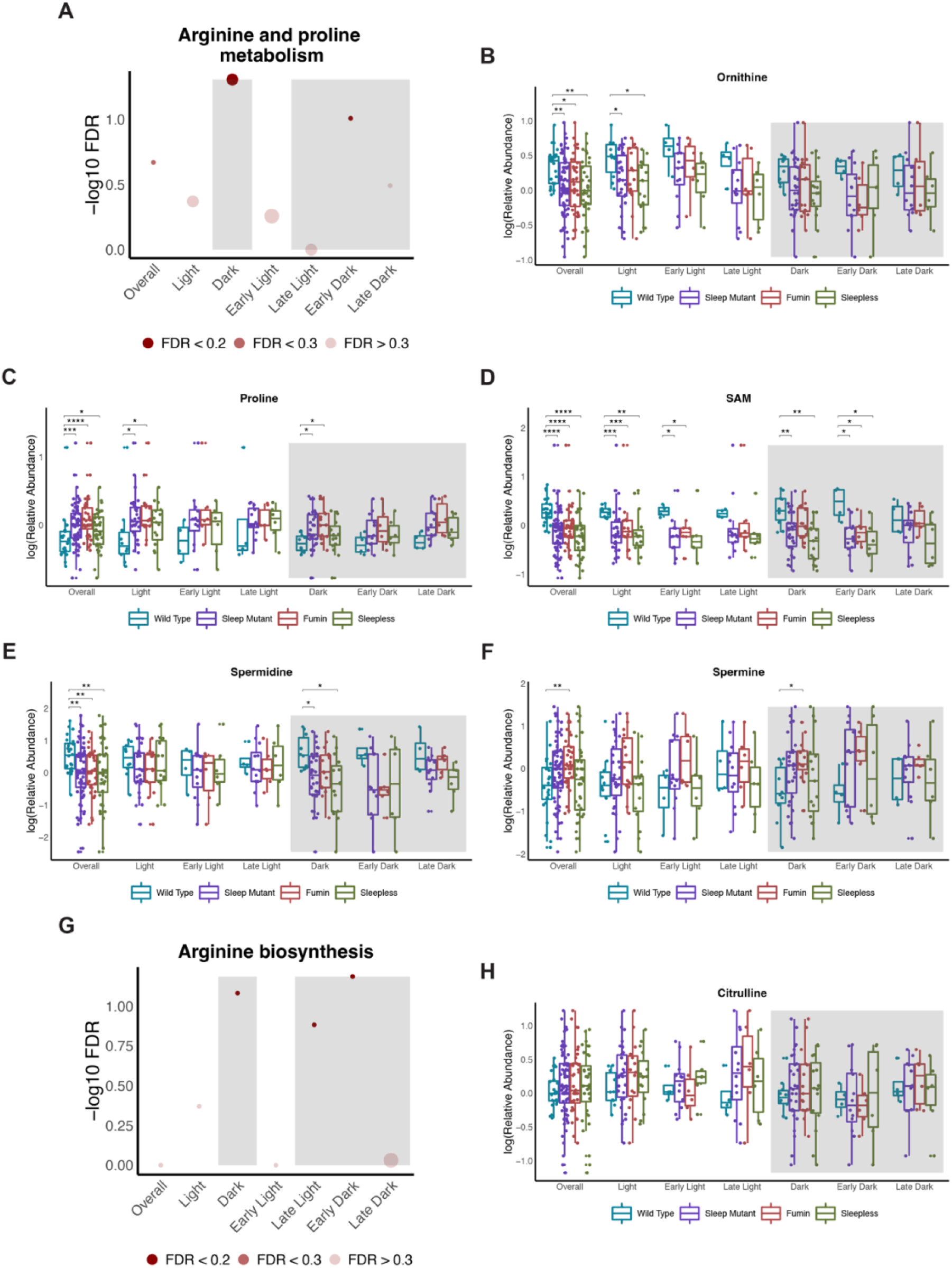
Arginine related metabolism is altered during the dark time period. (A and G) Negative log10 FDR values for arginine and proline metabolism (A) and arginine biosynthesis (G) pathways across the different significant OPLS-DA models. Shading represents FDR values. (B to F and H) Box plots of select metabolites associated with arginine and proline metabolism (B to F) and arginine biosynthesis (H) by groups (wild type, sleep mutants (consists of *fumin* and *sleepless)*, *fumin*, and *sleepless*) shown for different time periods. Significance was defined as a BH adjusted p-value less than 0.05 from a Wilcoxon test within each time period. Significance is indicated as follows: * < 0.05, ** < 0.01, *** < 0.001, **** < 0.0001.

### Alanine, aspartate, and glutamate metabolism is impacted during the early dark time points

Alanine, aspartate, and glutamate metabolism was one of the pathways significantly impacted between wild type and the short sleep mutants during the dark time points as well as during the early dark and late dark period (Figure 7A). This pathway did also trend towards being significantly impacted when time points were not considered. Interestingly, a number of the metabolites associated with this pathway such as alanine, aminosugar phosphates, carbamoyl phosphates, and GABA were significantly altered across the genotypes as well (Figures 7B, C, G and H). These metabolites were also altered during the dark time points, with alanine levels being decreased while aminosugar phosphates levels were elevated in the short sleep mutants both during the dark and early dark time points while carbamoyl phosphates and GABA were only significant during the overall dark period. Additionally, carbamoylaspartate was also significantly increased in the short sleep mutants during the dark period but was only significantly increased in the early dark period in *fmn,* although trends for increased levels in other groups compared to wild type could be observed (Figure 7F). Other metabolites associated with the pathway such as aspartate and asparagine were not significant at the univariate level, although trends could be observed for aspartate levels being increased in the short sleep mutants during the different dark periods and asparagine being decreased in the different groups during the early dark time points (Figures 7D and E). Overall, metabolites associated with alanine, aspartate, and glutamate metabolism were noted to be altered through univariate approaches during the dark time period.

**Figure 7.**
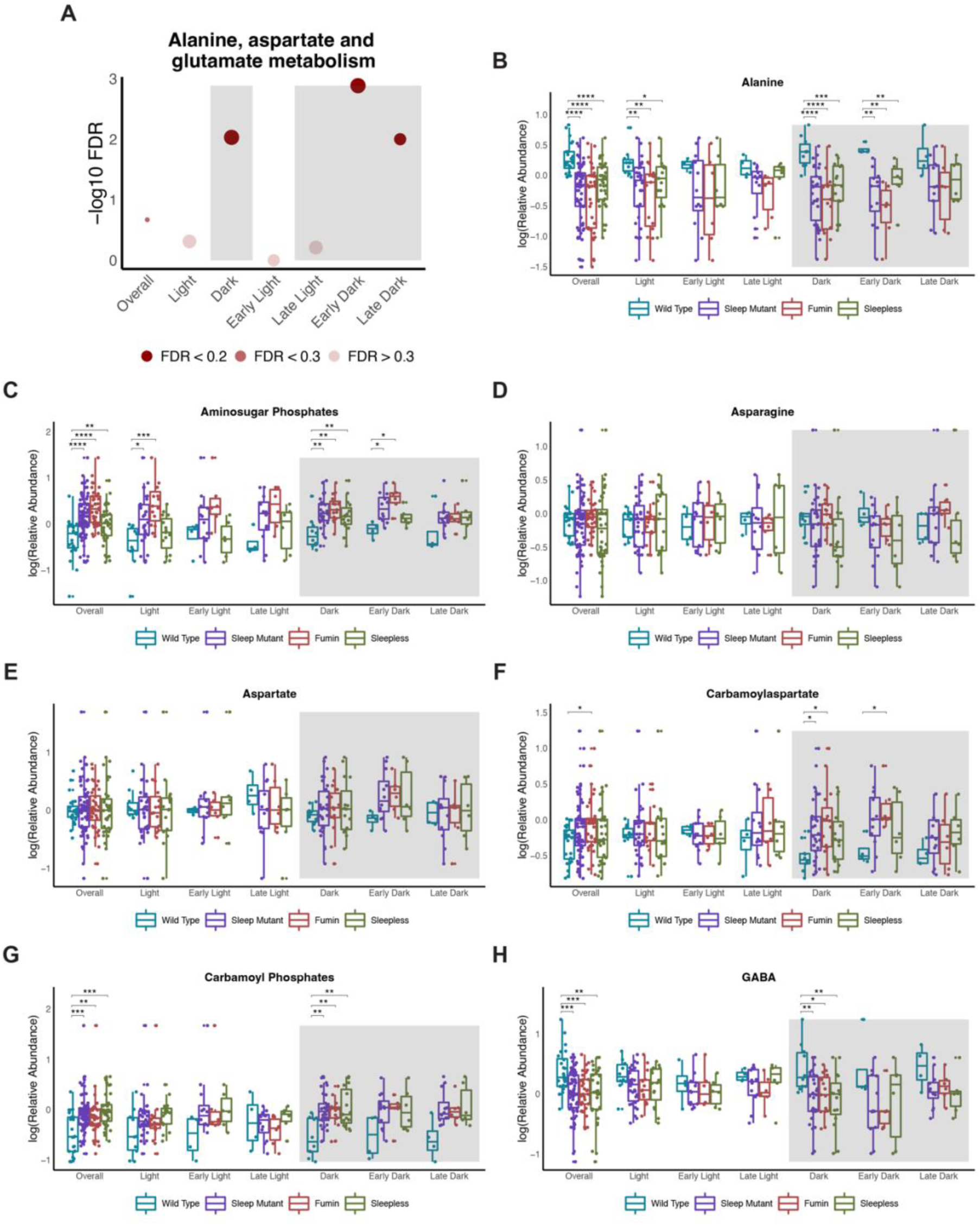
Alanine, aspartate, and glutamate metabolism is impacted during the early dark time points. (A) Negative log10 FDR values for alanine, aspartate, and glutamate metabolism pathway across the different significant OPLS-DA models. Shading represents FDR values. (B to H) Box plots of select metabolites associated with alanine, aspartate, and glutamate metabolism by groups (wild type, sleep mutants (consists of *fumin* and *sleepless)*, *fumin*, and *sleepless*) shown for different time periods. Significance was defined as a BH adjusted p-value less than 0.05 from a Wilcoxon test within each time period. Significance is indicated as follows: * < 0.05, ** < 0.01, *** < 0.001, **** < 0.0001.

### Altered glyoxylate and dicarboxylate and TCA cycle metabolism is observed in short sleep mutants

Two pathways, glyoxylate and dicarboxylate metabolism and citrate/TCA cycle, were significantly impacted during the late dark time points in pathway analyses using the identified metabolites from multivariate models (Figures 8A and B). Most of the metabolites identified as being associated with these pathways were common to both. For example, citrate/isocitrate, malate and pyruvate were common to both pathways and displayed significant differences overall between wild type and the short sleep mutants irrespective of time and during the overall dark period (Figures 8D, F and G). Interestingly, these metabolites in addition to aconitate and glutamine were not significantly different in the short sleep mutants overall from the wild type samples during the late dark period (Figures 8B to G). Although, trends in directionality were observed with decreased levels of aconitate, citrate/isocitrate and glutamine and increased levels of malate and pyruvate associated with the short sleep mutants during the late dark period. Overall, these results again highlighted differences in significant results between the multivariate and univariate approaches but indicated differences in these pathways to occur during the dark period.

**Figure 8.**
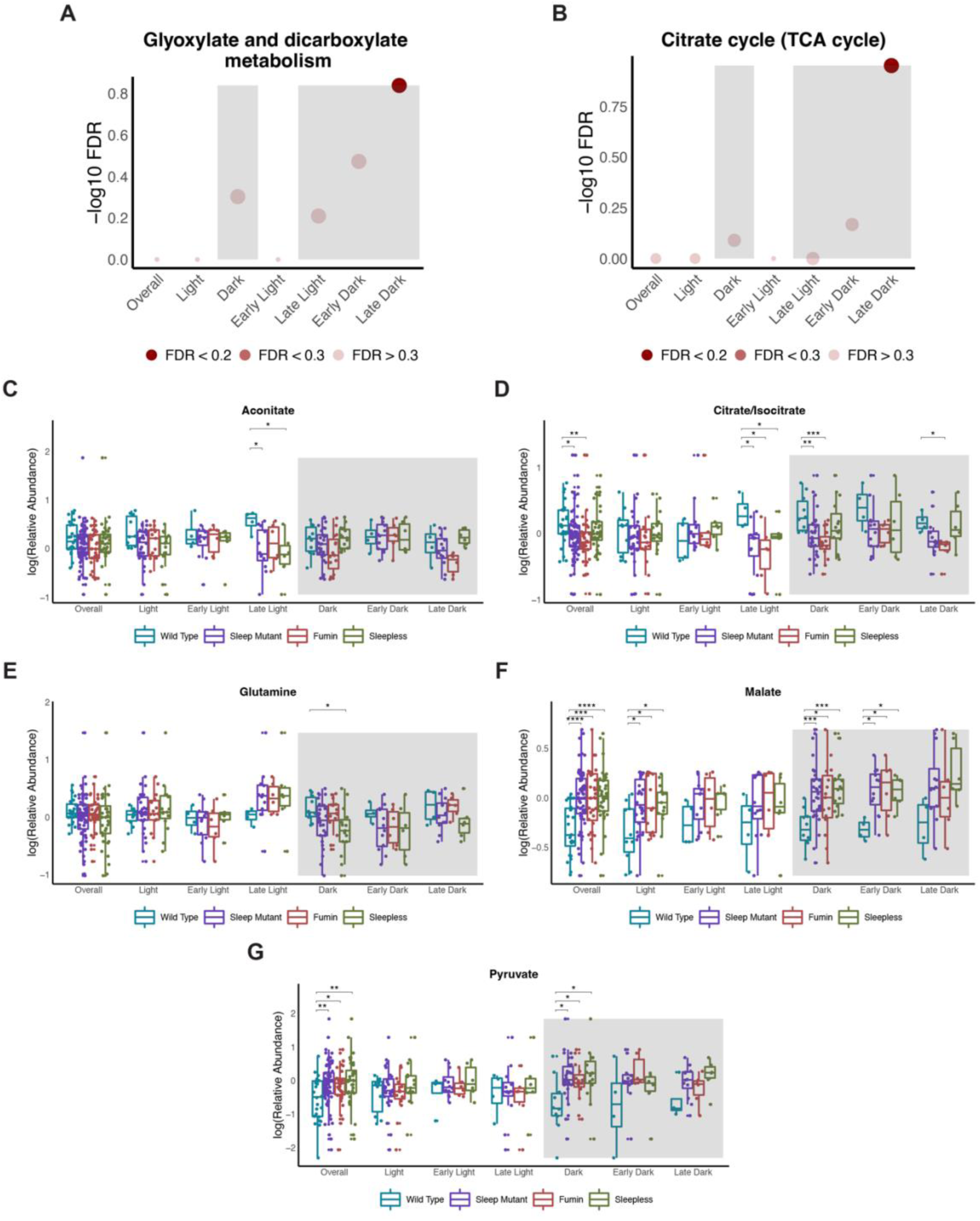
Altered glyoxylate and dicarboxylate and TCA cycle metabolism is observed in short sleep mutants. (A and B) Negative log10 FDR values for glyoxylate and dicarboxylate metabolism (A) and TCA cycle (B) pathways across the different significant OPLS-DA models. Shading represents FDR values. (C to G) Box plots of select metabolites associated with glyoxylate and dicarboxylate metabolism and the TCA cycle by groups (wild type, sleep mutants (consists of *fumin* and *sleepless)*, *fumin*, and *sleepless*) shown for different time periods. Significance was defined as a BH adjusted p-value less than 0.05 from a Wilcoxon test within each time period. Significance is indicated as follows: * < 0.05, ** < 0.01, *** < 0.001, **** < 0.0001.

### Short sleep mutants have altered purine and pyrimidine metabolism

Purine metabolism was a significantly enriched pathway during the late dark period (Figure 9 A). Similar to purine metabolism, many of the metabolites identified from multivariate models related to this pathway were observed to be significantly different from wild type irrespective of time. For example, metabolites such as adenosine were elevated while dAMP and hypoxanthine levels were decreased (Figure 9B to D). None of the metabolites were significantly different during the late dark period although trends in directionality were observed in similar directions to the earlier described changes. Significant changes for most of the metabolites were noted during the overall dark period though. For example, adenosine was significantly increased in the short sleep mutant while hypoxanthine was significantly decreased (Figure 9B and D). Another related pathway, pyrimidine metabolism, trended towards significance during the early dark period (Figure 9 E). Overall, most of the metabolites included in this pathway were also significantly altered in the short sleep mutants irrespective of time. For example, beta-alanine was decreased in the short sleep mutants compared to WT while dUMP and uridine were increased (Figure 9F to H). These metabolites also were significant with the same directionality during the overall dark and late dark period but not during the early dark period, but trends could be observed.

**Figure 9.**
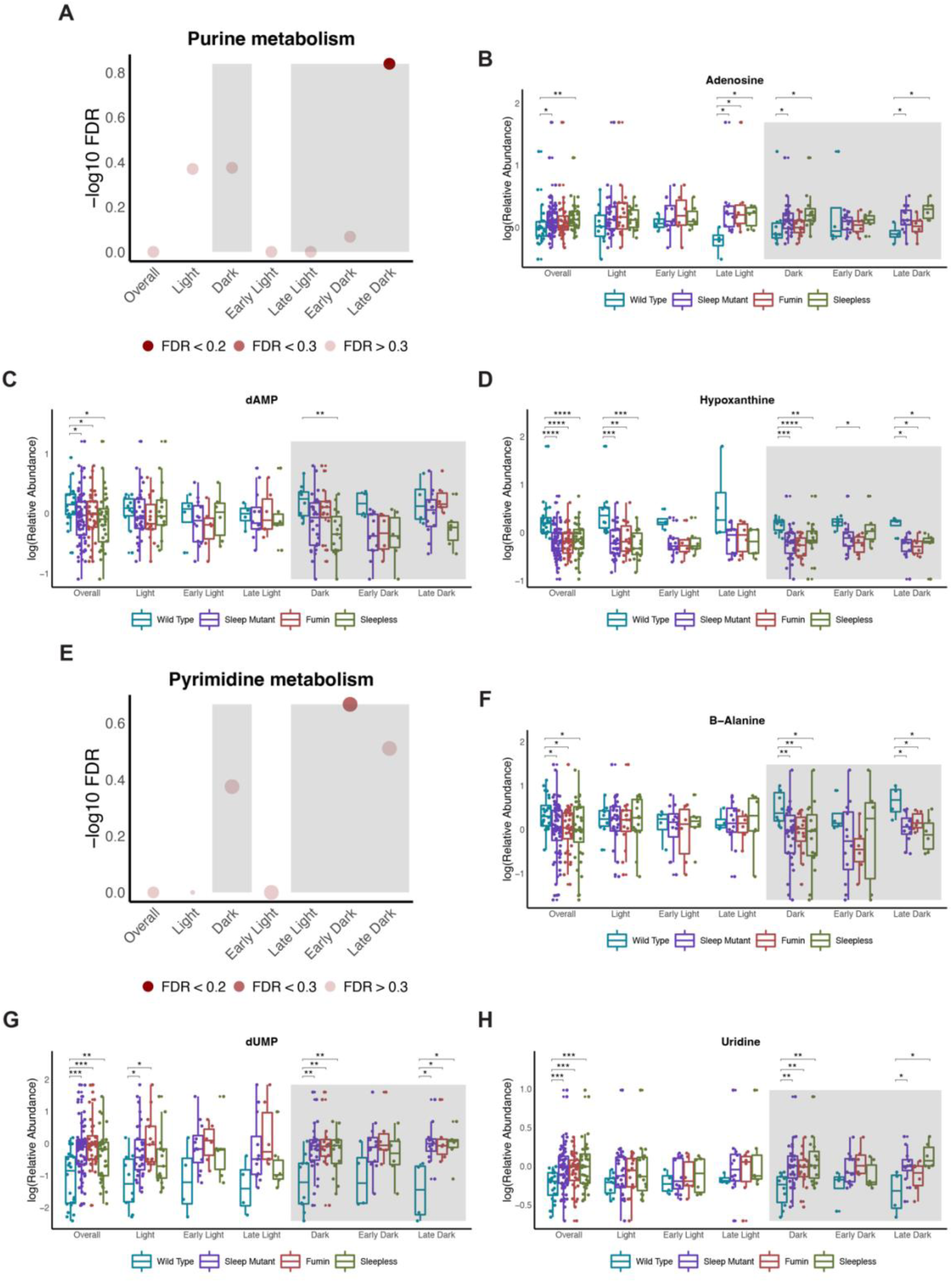
Short sleep mutants have altered purine and pyrimidine metabolism. (A and E) Negative log10 FDR values for purine metabolism (A) and pyrimidine metabolism (E) pathways across the different significant OPLS-DA models. Shading represents FDR values. (B to D and F to H) Box plots of select metabolites associated with purine metabolism (B to D) and pyrimidine metabolism (F to H) by groups (wild type, sleep mutants (consists of *fumin* and *sleepless)*, *fumin*, and *sleepless*) shown for different time periods. Significance was defined as a BH adjusted p-value less than 0.05 from a Wilcoxon test within each time period. Significance is indicated as follows: * < 0.05, ** < 0.01, *** < 0.001, **** < 0.0001.

### 24- and 12-hr rhythms are altered in short sleep mutants

Thus far, altered metabolites overall and according to time day had been noted in multivariate approaches in the short sleep mutants compared to wild type samples. To further determine whether the time-of-day differences were being driven by underlying changes in rhythmicity, we incorporated rhythmicity analyses. We tested for 24- and 12-hr rhythmic metabolites using both JTK and RAIN algorithms at different q-value cut-offs. Overall, the presence of 24-hr rhythmic metabolites was noted across all genotypes with the short sleep mutants having fewer rhythmic metabolites (Figure 10A). It is important to note that this trend although consistent between q-value cut-offs of 0.2 and 0.3 varied at q-value cut-offs of 0.1 and 0.4. However, the results at the q-value cut-offs of 0.1 and 0.4 may be impacted by the number of metabolites being detected. For example, more false positives may be identified at a q-value cut-off of 0.4. On the other hand, *fumin* had the largest number of 12-hr rhythmic metabolites followed by wild type and *sleepless* which had the lowest number of 12-hr rhythmic metabolites (Figure 10D). For further analyses, we used metabolites meeting a RAIN q-value cut-off of 0.2. Overall, the phases for the rhythmic metabolites were distributed across the day for metabolites with 24-hr periods however, *fmn* and *sss* displayed two major peaks (Figure 10 B). Furthermore, although both short sleep mutants had a peak for phases at ZT 16, the second peak occurred earlier in *fmn* at ZT 4 compared to ZT 6 in *sss.* For metabolites with a 12-hr period, overall similar phase distributions were noted across the different genotypes (Figure 10E). Next, to determine whether changes in rhythmicity may be driven by sleep, we compared the cycling metabolites between *fmn* and *sss.* For the 24-hr rhythmic metabolites, the presence of both unique and conserved (∼34% and ∼37% in *fmn* and *sss* respectively) rhythmic metabolites was noted (Figure 10C). On the other hand, a majority of the 12-hr rhythms were unique to *fmn* (Figure 10F). Furthermore, for both 24-hr and 12-hr rhythms, the common metabolites were not significantly enriched in any pathway (data not shown). Taken together, this highlighted that although conserved rhythmicity may be present between the short sleep mutants, each mutant is also uniquely impacted in terms of metabolite cycling.

**Figure 10.**
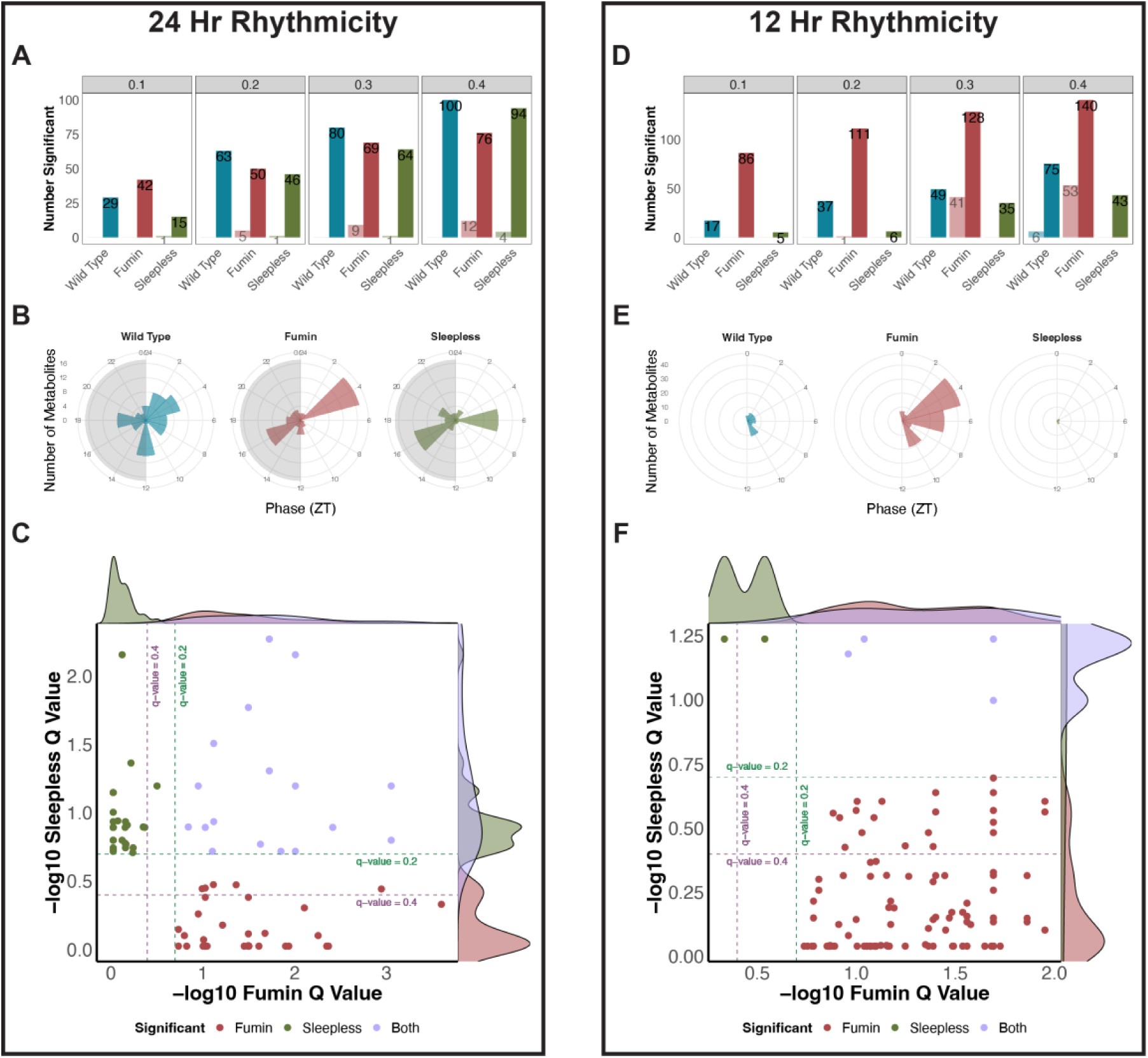
24- and 12-hr rhythms are altered in short sleep mutants. (A and D) Overview of the number of significant 24-hr (A) and 12-hr (D) rhythmic metabolites observed at RAIN (dark) or JTK (light) q-value cut-offs of 0.1, 0.2, 0.3, and 0.4 for each genotype. Colors represent genotype. (B and E) Phase profiles of 24-hr (B) and 12-hr (E) rhythmic metabolites at a RAIN q-value cutoff of 0.2. Colors represent genotype. (C and F) Comparison of 24-hr (C) and 12-hr (F) significantly cycling metabolites in *fumin* and *sleepless* defined by a RAIN q-value less than 0.2. q-value cutoffs of 0.2 (green dotted line) and 0.4 (purple dotted line) are shown. Colors refer to significant rhythmicity in one or both genotypes.

## Discussion

Disrupted sleep has been associated with various negative metabolic consequences. Most studies in this area have not been designed to study the time-of-day impacts of altered sleep. Here, we compared two *Drosophila* short sleep mutants with previously acquired wild type samples where all genotypes were collected at 2-hr time resolution, thus enabling time-of-day changes to be examined. Overall, we were able to identify a common metabolic signature to both short sleep mutants potentially arising from decreased sleep. Impacted metabolites were noted to be associated with nicotinate and nicotinamide metabolism. Moreover, dark periods were found to be associated with an increased number of altered metabolic pathways. As *Drosophila* are diurnal, sleep disruption is most likely to occur during the dark period as well. Taken together, this suggested that decreased sleep caused a more noticeable impact during periods where sleep would normally occur. Unsurprisingly, although similar trends could be observed in metabolites involved in the significantly impacted pathways, not all were significant through univariate analyses. However, this is not unexpected as both multivariate and univariate methods approach the data in different ways. Multivariate analyses provide insight into overall and/or concerted changes across different metabolites whereas univariate tests look at each individual metabolite separately. Therefore, even though not all metabolite-level differences were significant at a univariate level, they were still significantly contributing to the separation of short sleep mutants from wild type samples since they were identified as VIP metabolites in the discriminant models.

### Altered nicotinate and nicotinamide metabolism suggests decreased NAD+ levels

One of the pathways significantly impacted in short sleep mutants overall irrespective of time in addition to time-dependent differences was nicotinate and nicotinamide metabolism. Nicotinate and nicotinamide metabolism has also been previously reported to be impacted by decreased sleep. For example, in a 7-day paradoxical sleep deprivation study, it was a significantly impacted pathway [22]. Two of the metabolites measured in the study were also identified as VIP metabolites in our results. Nicotinamide levels were elevated in the short sleep mutants and consistent with the observation in the paradoxical sleep deprivation group while nicotinic acid levels were increased in the short sleep mutants while they had been previously reported to be decreased under paradoxical sleep deprivation. Moreover, a greater number of metabolites involved in the nicotinate and nicotinamide metabolism pathway were identified in the short sleep mutants. Specifically, in addition to increased levels of nicotinic acid and nicotinamide, decreased levels of NAD+, NADP+, nicotinamide riboside, and nicotinamide ribotide were noted during each time period tested (overall, light, dark, early/late light and dark). Overall, this potentially pointed towards increased NAD+ utilization and/or decreased NAD+ biosynthesis.

NAD+ has been identified in circadian studies as well. For example, NAD+ levels have been observed to decrease with age [23, 24]. Here, supplementation with nicotinamide riboside was noted to prevent the decline of circadian function in aging mice [23, 24]. Since nicotinamide riboside was a precursor to NAD+, this pointed to a potential role for NAD+ in preventing age-related decline of circadian function [23, 24]. Additionally, NAD is known to modulate the DNA binding affinity of *Clock* and *BMAL1* with circadian transcription factors. Specifically, the reduced forms, NADH and NADPH, enhance DNA binding while the oxidized forms, NAD and NADP, inhibit binding [25]. SIRT1, a NAD+ dependent protein deacetylase, has also been observed to have circadian properties and to influence rhythmicity of core clock genes [26, 27]. Taken together, decreased levels of several metabolites in the nicotinate and nicotinamide metabolism pathway may lead to decreased levels of NAD+ in the short sleeping mutants. Decreased NAD+ levels may implicate underlying circadian dysfunction in the short sleep mutants and potentially a premature aging phenotype.

### Altered arginine related metabolism may be driven by circadian processes

Two pathways related to arginine metabolism, arginine and proline metabolism and arginine biosynthesis, were noted to be impacted during the dark period. Arginine and proline metabolism also was a pathway likely to be impacted overall in the short sleep mutants as well as during the early dark period while arginine biosynthesis was also significantly impacted during the late light and early dark periods. Recently, metabolites involved in nitrogen and polyamine metabolism related metabolites, some of which overlap with arginine related metabolism, were identified to be impacted in heads of *Drosophila* short sleep mutants (*fmn, sss,* and *rye*) [26]. For example, increased levels of spermidine and decreased levels of GABA were noted in sleep mutants across both studies. Interestingly a number of metabolites showed opposite changes in our analysis of fly bodies versus what had been reported for fly heads. For example, proline was elevated in the short sleep mutants but was noted to be decreased in the heads, while ornithine and acetylputrescine levels were decreased in the short sleep mutant bodies but were elevated in the heads. Additionally, some metabolites such as citrulline and arginine were not significantly altered in fly heads; however, in the measurements of fly bodies, we observed decreased levels of arginine during the dark period and increased citrulline levels especially during the light period. Although two of the same short sleep mutants were used in the study, the differences in the directionality of changes observed in our study can be explained by a few differences in study design. One potential reason can be differences in metabolism in head versus fly body. Additionally, the metabolite measurements in the fly heads were performed in fly samples collected at a single time point (ZT 6) whereas our study included samples collected across a day. This does highlight the importance of looking at specific regions and/or tissues to discern the impact of sleep but also of looking at differences across time of day. Previously, arginine and proline metabolism was also noted to be altered in insomnia patients during the day and night time periods [10]. This is similar to what we observed in the sleep mutants with the pathway being significant during both the late light and early dark period. Moreover, arginine and proline metabolism has also been observed to be a significant circadian pathway in mice and was a significantly impacted pathway under constant routine conditions in humans irrespective of the day or night-shift groups [28, 29]. Overall, the significant changes observed in arginine and proline metabolism may also be driven by underlying circadian processes.

### Alanine, aspartate, and glutamate metabolism may be impacted uniquely in response to sleep loss

Short sleep mutants also had altered alanine, aspartate and glutamate metabolism during the dark period including early and late dark time points. In a chronic sleep fragmentation study, metabolites associated with alanine, aspartate and glutamate metabolism were altered. For example, aspartate was significantly decreased in response to 15 days of sleep fragmentation [30]. In the short sleep mutants, however, these metabolites were increased during the dark period. The differences in the two results may be due to the different metabolic impacts of sleep fragmentation versus sleep loss.

### Glyoxylate and dicarboxylate and TCA cycle metabolism are altered as a result of common metabolites

Glyoxylate and dicarboxylate and TCA cycle metabolism was another significantly impacted pathway in a paradoxical sleep deprivation study. Although none of the VIP metabolites associated with glyoxylate and dicarboxylate metabolism in the short sleep mutants overlapped with the paradoxical sleep deprivation study, overall, the pathway was noted to be decreased in both. In the short sleep mutants, three of the four metabolites associated with the pathway had decreased levels during the dark period (aconitate, citrate/isocitrate, and glutamine) [22]. Similarly, another study noted decreased levels of oxalic acid, another metabolite associated with glyoxylate and dicarboxylate metabolism, after sleep restriction; however, levels returned to normal after recovery sleep in both humans and rats [31].

Alternatively, three of the four metabolites identified as being involved in glyoxylate and dicarboxylate metabolism can also be involved in TCA cycle metabolism. Altered TCA cycle metabolism has been reported previously as a result of chronic intermittent hypoxia in mice [32]. In a more recent sleep study, TCA cycle intermediates were noted to be associated with sleep-wake states with elevated levels observed during the transition to REM (rapid eye movement) sleep [33]. Metabolites related to TCA cycle metabolism were noted to be rhythmic in U2OS cells [34]. As most of the metabolites (pyruvate, malate, citrate/isocitrate, and aconitate) are more closely related to TCA cycle metabolism, it is possible that this is the pathway impacted. However, this provides an example of limited insight into how metabolite levels are arising and would necessitate the use of labelled tracers to determine whether TCA cycle or glyoxylate and dicarboxylate metabolism are impacted.

### Purine and pyrimidine metabolism

Short sleep mutants had significant differences in purine metabolism during the late dark period and were likely to have altered pyrimidine metabolism during the early dark period. Both purine and pyrimidine metabolism play an important role in energy-related metabolism and can serve as precursors for nucleotide biosynthesis and nucleotide cofactors [35]. Purine metabolism has previously been reported as being altered under a 14-day chronic sleep disruption protocol while increased pyrimidine metabolism has been associated with the wake period [36, 37]. Both metabolic pathways have also been impacted in circadian studies. For example, dampened cycling of purine catabolism has been observed in response to a high fat diet while rhythmicity of both purine and pyrimidine metabolism has been noted in zebrafish, another model for sleep and circadian studies [38, 39]. Both of these pathways may be influenced by both circadian and sleep driven changes. As sleep mutants, may have an increased energetic demand, these pathways may be more active during the dark period.

### Neurotransmitters are impacted by decreased sleep

Several neurotransmitters were observed to be VIP metabolites across different models. For example, GABA was decreased in the short sleep mutants in comparison to wild type whereas acetylcholine and adenosine were elevated in the short sleep mutants. Glutamate was elevated overall and during the light period but showed some variability during the dark period. Neurotransmitters have also been known to play a role in regulating sleep. For example, acetylcholine promotes wakefulness while GABA and adenosine promote sleep [40]. Increased acetylcholine and decreased GABA were consistent with the roles of these transmitters in promoting wakefulness and promoting sleep, respectively. Adenosine however is elevated and may indicate increased sleep need in the short sleep mutants.

Overall, we have been able to identify common metabolic profiles associated with decreased sleep in two short sleep mutants. Additionally, time of day variability also is present across the genotypes both in terms of rhythmicity and at a metabolome-level. During the dark period, several metabolic pathways are impacted in the short sleep mutants. However, several metabolites are involved in multiple pathways and so the question remains whether they are involved in one or both of the pathways. For example, TCA cycle and glyoxylate and dicarboxylate metabolism contained almost identical metabolites and would require the use of additional experiments such as those incorporating stable isotopes to determine which of the two pathway(s) was altered.

## Methods

### Drosophila Strains

Two *Drosophila melanogaster* short sleep mutants, *fumin* [79] and *sleepless* [80], were maintained on standard cornmeal/molasses medium at 25°C under 12:12 LD conditions.

### Time Course Collection

Eclosed male flies were collected daily for 3 days. On day 4, 15 male flies were sorted into new vials for each time point (ages were evenly distributed among all time points) and were entrained into appropriate LD cycles for a minimum of 3 days. For *sleepless,* due to limited number of age-matched flies, 24-hr collections were performed across two days with ZT 0, 4, 8, 12, 16, 20 collected on one day and ZT 2, 6, 10, 14, 18, 22 collected on a second day. All flies were between 5 to 7 days old at the time of collection. Flies were anesthetized with carbon dioxide, collected on dry ice, and stored at -80°C until metabolite extractions. Three separate days of collections (biological replicates) were performed for each sleep mutant.

### Metabolite Extraction and LCMS Measurements

Twelve fly heads and bodies were separated on ice for each sample and kept at -80°C. A modified Bligh-Dyer extraction was performed on fly bodies [41, 42]. Briefly, a stainless-steel bead and 600 μL of 2:1 cold methanol:chloroform was added to each sample followed by homogenization for a total of 4 minutes at 25 Hz in a tissue homogenizer. Next, 200 μL each of milliQ water and chloroform were added to each sample. Samples were vortexed and centrifuged for 7 minutes at 18787xg at 4°C. From each sample, 350 μL of the upper fraction containing the polar metabolites was collected and dried under vacuum for 6 hours. Dried polar metabolites were resuspended in 100 ul of acetonitrile:milliQ water and vortexed for 20 seconds. Samples were analyzed using an ion-switching LCMS method using instrumentation, MRM transitions, chromatography, mobile phases, and parameters described previously in [41]. For each sample, two 5 ul injections (analytical replicates) were analyzed. A quality control sample comprising a pool of all samples being analyzed was also run at the beginning and end of the run as well as every 10th injection.

### Data Processing and Normalizations

Along with a previously acquired dataset for wild type [21], all chromatograms were processed using El-MAVEN (version 0.12.1 beta) to obtain ion counts. Ion counts were used for further processing in R on both data sets (wild type and short sleep mutants) separately. Any sample displaying a large column pressure fluctuation at the start of a chromatographic run was excluded from further analyses due to improper injection. A LOESS correction using the quality control samples was performed to correct for instrument drift [43]. Additional criteria included removing metabolite features displaying a relative standard deviation greater than 30 percent or if a metabolite was missing in more than 50 percent of the QC samples. PCA (described in Section 2.5.5) was performed at this step to assess data quality. Clustering of QC samples was observed in an initial PCA; however, improved distribution of loadings and scores were observed with an additional median fold change normalization step. Therefore, the median fold change normalized data was combined across both data sets. To normalize for batch differences, a ratio of each metabolite measurement was taken to the measurement at ZT 0 on a group- and biological replicate-level. Any metabolite that was not detected in at least half of the samples for each batch was removed from the overall dataset. The remaining metabolites were used for further analyses.

### Multivariate and Rhythmicity Analyses

Multivariate analyses were performed using SIMCA (Umetrics, version 17). Principal component analyses were performed on raw, instrument drift corrected, and normalized data to determine the optimal data set to use based on the absences/presence of outliers, clustering of quality control samples and analytical replicates, and overall distribution of scores and loadings. The combined ratio data for all groups was further used in orthogonal partial least squares-discriminant analyses to determine group- and time-level differences. In order to determine similarities between different OPLS-DA models, a shared and unique structures (SUS) approach was also used.

To assess rhythmicity, JTK (R MetaCycle package) and RAIN (R Rain package) algorithms were used to test for 20-28-, 24-, 12-, and 8-hr cycling.

### Pathway Analysis

Identified VIP metabolites and/or significant metabolites from univariate tests were used in metabolic pathway analyses through MetaboAnalyst 5.0 [44]. Metabolites were uploaded using HMDB identifiers and processed using a hypergeometric enrichment method, relative-betweenness centrality for topology analysis using the *Drosophila melanogaster* (KEGG) pathway library. The resulting data was imported into R (Version 4.0.3) and further analyzed. Significant pathways were defined using a FDR less than 0.2 (with those less 0.3 identified as potentially impacted pathways). To be included in analyses, an impact value greater than zero was required for at least one time comparison.

## Acknowledgements

We thank Lisa Bottalico for discussions on LCMS methodology; Gregory Grant for discussions on considerations for proper sample randomization; Tom Brooks and Antonijo Mrcela for discussions on cycling measurement algorithms; and Joeseph Bedont for discussions around short sleeping mutants. This work was supported by NIDDK of the National Institutes of Health under award number R01-DK120757. D.M.M. was supported through a Pharmacology T32 Training Grant (T32 GM008076). A.S. is an investigator of the Howard Hughes medical Institute (HHMI).

## Author Contributions

Conceptualization, D.M.M., A.S., and A.M.W.; Methodology, D.M.M., A.S.G., and A.M.W.; Formal Analysis D.M.M., and A.M.W.; Investigation, D.M.M., and A.S.; Resources, A.S., and A.M.W.; Writing – Original Draft, D.M.M., and A.M.W.; Writing – Review & Editing, D.M.M., A.S., A.M.W; Visualization, D.M.M, A.S, A.M.W.; Supervision A.M.W.; Funding Acquisition, A.S. and A.M.W.

